# Comment on Smaers *et al.* (2016): A nonviable phylogenetic comparative method hampered by circularity, inaccuracy, and bias

**DOI:** 10.1101/058420

**Authors:** Randi H. Griffin, Gabriel S. Yapuncich

## Abstract

Smaers, Mongle & Kandler (2016) (*Biological Journal of the Linnean Society*, 118: 78-94) introduced a new phylogenetic comparative method, multiple variance Brownian motion (mvBM), for reconstructing ancestral states given a phylogenetic tree and continuous trait data. The authors conducted a simulation study and argued that mvBM outperforms constant variance Brownian motion (BM) when rates of evolution vary across the phylogeny. In this comment, we argue that mvBM is not a viable statistical method because it is fundamentally a circular analysis that overfits phylogenetic branch lengths to the data. We further argue that the comparison of mvBM to BM under conditions where the assumptions of BM are clearly violated is not an informative performance analysis, and that the simulation study of Smaers *et al*. (2016) exaggerates the performance of mvBM by focusing on a narrow range of simulation conditions and reporting aggregated accuracy metrics that obscure severe inaccuracy and bias in its ancestral state estimates. Our arguments are supported by simulation results. We conclude that mvBM is not a viable phylogenetic comparative method.

## Introduction

Smaers, Mongle & Kandler (2016) introduced a new phylogenetic comparative method (PCM), which they call multiple-variance Brownian motion (mvBM), designed to estimate ancestral states from a phylogeny and continuous trait data. The authors validated their new method with simulations showing mvBM produces similar or improved ancestral state reconstructions (ASRs) compared to constant variance Brownian motion (BM). The mvBM method is partially based on the Independent Evolution (IE) method introduced by Smaers & Vinicius (2009).

We recently demonstrated serious theoretical and statistical problems with IE (Griffin & Yapuncich, 2015), leading us to examine mvBM closely. Here, we focus on two aspects of Smaers et al’s (2016) simulation study that deserve further examination. First, we explore why mvBM appears to produce likelihoods that are substantially greater than those of BM when data are simulated with a BM model, even though Smaers *et al.* assert that mvBM is identical to BM when BM is the true model. Second, we investigate the accuracy of mvBM ASRs at specific nodes when data are simulated under the “burst of evolution” scenario (i.e., one or more phylogenetic branches have increased evolutionary rates), which mvBM is supposedly designed for. Smaers *et al.* did not report accuracy measures for individual nodes in their simulation study; rather, they aggregated accuracy measures across all nodes in the tree.

In this comment, we argue that mvBM is a nonviable PCM and that Smaers *et al.* conducted an inadequate simulation study. Specifically, we argue: 1) mvBM is a *circular analysis* that overfits phylogenetic branch lengths to the data, 2) mvBM has improved accuracy compared to BM under the burst scenario due to similarities between Smaers *et al.*’s method and phylogenetically independent contrasts (PIC; Felsenstein, 1985), and 3) the aggregated accuracy metrics presented by Smaers *et al.* mask extreme bias and inaccuracy in mvBM’s ASRs. Simulations were performed in R (R Core Team, 2015) with the packages ‘ape’ (Paradis *et al,* 2004), ‘phytools’ (Revell, 2012), and ‘geiger’ (Harmon *et al,* 2008). Supplementary Information S1 contains a detailed description of our implementation of mvBM. Code for replicating our simulations is available on GitHub (Griffin, 2016).

### mvBM is a circular analysis that overfits phylogenetic branch lengths to the data

The mvBM method estimates ancestral states in four steps: 1) “the phenotypic trait values of the *n* – 1 internal nodes” (p. 80) are estimated using a phylogeny and trait data for terminal taxa, 2) these values are used to estimate squared change along each branch, 3) each branch is then elongated by an amount proportional to its inferred squared change, and 4) “the trait values for all internal nodes” (p. 82) are estimated by fitting a standard BM model to the mvBM-transformed phylogeny. Smaers *et al.* say “the [equations] used for estimating the nodal values can be modified” (p. 81) and “alternative branch lengthening procedures can be conceived” (p. 82), thus the specific equations used in steps 1 and 3 are not essential features of mvBM. Rather, the key feature distinguishing mvBM from other ASR methods is the repeated estimation of internal node values in steps 1 and 4. In their description, Smaers *et al.* distinguish between the first and final sets of estimated “phenotypic values at internal nodes” by suggesting that the first set is used to “parameterize a mvBM model” (p. 80), while the second set represents ASRs “under a multiple variance BM model of phenotypic evolution” (p. 82). However, it is difficult to interpret the first set of “phenotypic values at internal nodes” as anything other than ASRs, and Smaers *et al.* give no indication as to what else they might be. The first set of nodal values are used to infer change along individual branches as the difference between ancestral and descendent nodal values, which only makes sense if the nodal values represent ASRs for the trait in question. Because the starting and ending points of the algorithm are the same (i.e., estimated trait values at internal nodes), mvBM is fundamentally a circular analysis.

This circularity explains why the likelihood of data under a BM model improves when the original phylogeny is replaced with the mvBM-transformed phylogeny, even when data were simulated under BM (Smaers et al., 2016, their Figure 3). The only way a model can consistently fit data better than the generating model is by *overfitting,* i.e., modeling random noise as meaningful signal (McElreath, 2016). The mvBM method overfits by elongating phylogenetic branches such that larger changes are made more likely on branches where larger changes have been inferred from the initial ASRs. Unsurprisingly, the likelihood of the data improves when this mvBM-transformed phylogeny is used to represent the data’s covariance structure in a BM model. This approach presents a major problem for statistical inference: even when evolution follows a single-rate BM process, mvBM *always* estimates 2*N-* 2 rates and produces an altered phylogeny that fits the data better than the original phylogeny. Since this altered phylogeny largely captures random noise, it provides no meaningful insights about the data. Moreover, the claim that mvBM estimates nearly twice as many rates as there are observations raises red flags: no statistical method can estimate more independent parameters than there are observations (see Griffin & Yapuncich 2015 for a discussion of this problem in relation to the IE method of Smaers & Vinicius 2009). This is why multiple-regime models of evolution include mechanisms to avoid overfitting, such as penalized likelihoods, Bayesian priors, or caps on the number of parameters (reviewed in Ho and Ané 2014), but mvBM employs no such mechanisms. Simulations demonstrating mvBM’s overfitting are provided in Supplementary Material S2.

Before discussing Smaers et al.’s simulation study, we must emphasize that mvBM represents a major departure from the basic principles underlying modern PCMs. The statistical terminology of Smaers *et al*. suggests their method relates to existing statistical models of evolution, particularly the multiple-rate BM model of Venditti, Meade & Pagel (2011). However, a true statistical model of evolution can be represented by a likelihood function describing the mathematical relationships among parameters (*sensu* Venditti *et al.* 2011), permitting parameter optimization and formal model comparison (e.g., using likelihood ratios or AIC). The mvBM method includes no likelihood function, therefore it cannot be used for parameter optimization or formal model comparison, nor can it be accurately described as a “multiple-rate BM model” (which implies a parametric model of evolution that extends the BM model). Although Smaers *et al.* confusingly report likelihoods for mvBM, these are invalid likelihoods as they do not account for the 2*N*’ 2 branch-specific rates estimated in steps 1-3. Instead, these likelihoods correspond to a BM model with two parameters (mean and variance) fit to the original data, with the overfit mvBM-transformed phylogeny incorporated into the covariance structure. These likelihoods are useless for statistical inference.

*How does mvBM produce more accurate ASRs than BM under a “burst” scenario?*

Smaers *et al.* (2016) simulated data under a “burst” scenario, where the rate of BM evolution is increased by 2-4 orders of magnitude on one or five branches of the phylogeny. They demonstrated that mvBM produces more accurate ASRs relative to BM when data are generated under the burst scenario. In this section, we show that in practice, mvBM produces ASRs that are roughly intermediate between BM (i.e., global maximum likelihood) and phylogenetically independent contrast (PIC) ASRs (i.e., local maximum likelihood; Maddison 1991). We argue that the improved accuracy of mvBM relative to BM under the burst scenario is a result of its similarity to PIC ASRs.

In step 1 of mvBM, Smaers *et al.* traverse the tree from the tips to the root in the same fashion as the PIC algorithm introduced by Felsenstein (1985). At each node, the ASR is computed as a weighted average of a “global” and “local” estimate for the trait value at that node. The “global” estimate is a weighted mean of all the tips of the phylogeny, with the tips weighted by their inverse squared patristic distance to the node in question (Smaers et al.,2016: Eq. 2). The “local” estimate is the average value of the immediate descendants of the node in question, without accounting for phylogenetic distance (Smaers et al.,2016: Eq. 3). Finally, the “global” and “local” estimates are averaged (Smaers et al,2016: Eqs. 4 and 5), with the “local” estimate being given twice the weight of the “global” estimate (this is arbitrary, and Smaers *et al.* indicate different weighting schemes are allowed).

This approach is not justified based on any current theoretical models of evolution. The “global” ASR is nearly identical to the problematic “adaptive peak” equation of Smaers & Vinicius (2009), although branch lengths are now squared. We have previously argued that it makes no theoretical sense to estimate an “adaptive peak” with this equation (Griffin & Yapuncich, 2015), and we see no theoretical justification why this particular value should now be considered a “global” estimate of an ancestral state. We also note that the “local” ASR is non-phylogenetic since it ignores branch lengths. While these criticisms speak to the theoretical weakness of the approach, we focus here on the fact that it produces similar results to the X’s computed with Felsenstein (1985) PIC algorithm. Maddison (1991) showed that PIC ASRs are identical to the local maximum likelihood ASR at each node. Since Smaers *et al.* square the inverse patristic distances in their “global” estimate, and the “local” estimate is weighted more heavily, the influence of phylogenetically distant tips is reduced, bringing the initial mvBM ASRs closer to PIC ASRs. These initial ASRs are then used to infer branch-specific rates of change, elongate branches according to the inferred change on the branch, and estimate BM ASRs on the altered tree. This essentially weights branches such that the initial ASRs become more likely under a BM model. As a consequence, when BM ASRs are estimated for a second time on the altered tree, the results are biased toward the initial ASRs, which are similar to PIC ASRs.

The similarity between PIC ASRs and the ASRs computed in step 1 of mvBM explains why mvBM produces more accurate ASRs than BM under a “burst” scenario. The ASRs of BM are truly “global” in the sense that they represent the set of ASRs with maximum likelihood under a BM model. As such, when a large burst occurs on a single branch, BM spreads the burst across multiple branches and affects ASRs for all internal nodes. In contrast, PIC produces “local” ASRs that represent the maximum likelihood ASRs when only the focal node’s immediate descendants are considered. Consequently, a burst of evolution will have a more local impact on PIC ASRs.

To illustrate this point, we performed a simple set of simulations. We generated a phylogeny with 3 tips, such that there is one internal node besides the root (Figure 1a). We simulated 100 continuous traits with mean 0 and variance 0.01 along the branches leading to the pair of sister taxa (branches 1-3), but systematically varied trait change on the long branch (branch 4) from 0 to 1.5, representing evolutionary bursts of different magnitudes. We then used maximum likelihood BM, mvBM, and PIC to reconstruct the ancestral state at the internal node (node 5) for each simulated dataset, and compared the results. Figure 1b shows that BM and mvBM become increasingly inaccurate as the magnitude of the burst on branch 4 increases, while the accuracy of PIC is unaffected. Since PIC computes local estimates of the ancestral states, the estimate at node 5 is unaffected by the burst on branch 4. In contrast, BM spreads the burst across branches such that large positive changes on branch 4 lead the ASR at node 5 to be overestimated. Since mvBM is neither entirely local nor entirely global, in this particular context it produces ASRs of intermediate accuracy.

**Figure 1:**
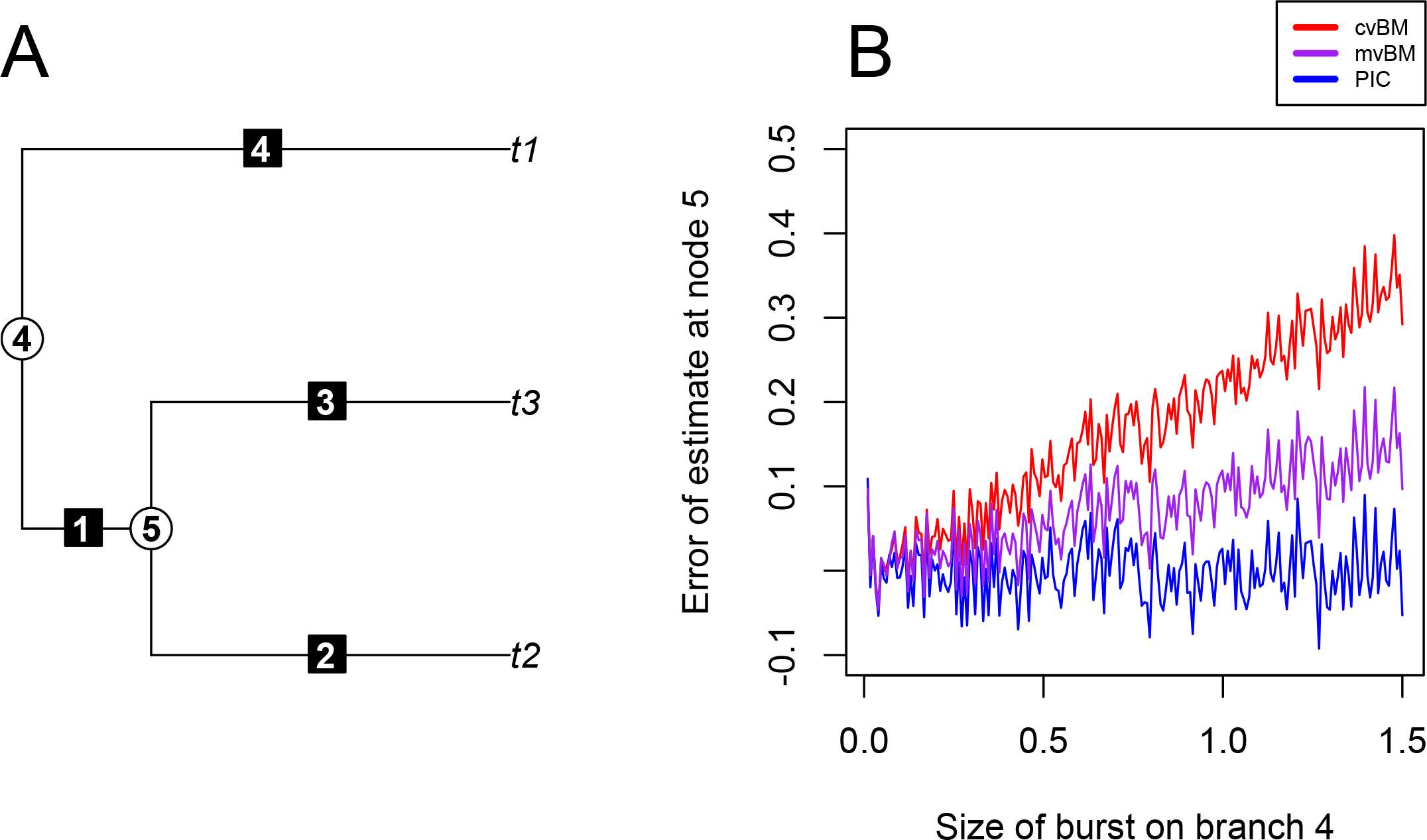
A) The phylogeny used for simulations demonstrating error and bias in mvBM ancestral state reconstructions. A Brownian motion model is used to simulate 100 traits with variance 0.01 on branches 1-3, while the trait change on branch 4 is varied from 0 to 1.5. For each simulation, the ancestral state at node 5 is estimated using PIC, mvBM, and BM. B) Relationship between different magnitudes of trait change on branch 4 and overestimation of the ancestral state at node 5 based on PIC, mvBM, and BM.

Of course, this does not mean that BM is a bad model or that PIC is better. Rather, BM is appropriate when BM is a reasonable model for the data, which it clearly is not when evolutionary rates are increased by orders of magnitude on individual branches. Importantly, Smaers *et al.*’s demonstration that mvBM performs relatively better than BM under simulation conditions where BM is clearly inappropriate does not constitute a rigorous performance analysis. Rather, a rigorous performance analysis must demonstrate that a method accurately estimates the parameters it is intended to estimate. Below, we present such a performance analysis and demonstrate severe bias and error in mvBM ASRs under the “burst” scenario it is purportedly designed for.

### Bias and error in mvBM ASRs

Using a 100-taxa phylogeny, Smaers *et al.* simulated 1000 datasets with 1) a burst on one branch, 2) five bursts spread across the tree, and 3) five bursts clustered together. The rate of evolution was increased on “burst” branches by either 100, 2500, or 10,000 times the baseline rate of evolution. For each simulated dataset, they estimated the *R*^2^ of a linear regression of mvBM-estimated ancestral states against true ancestral states. They found the vast majority of *R*^2^ values were >0.95 (Smaers et al.,2016, their Figures 4-6), suggesting a close fit between mvBM estimates and true ancestral states. They concluded that mvBM produces accurate ASRs even when there are large bursts of evolution on individual branches of the tree. However, Smaers *et al*. did not report ASRs for specific nodes near the bursts, nor did they vary the location of the bursts in the phylogeny. Here, we simulate the single-burst scenario to show that 1) error in ASRs at the base of the “burst branch” is large and biased, 2) by simulating a large number of tips relative to bursts and aggregating results across all nodes, Smaers *et al.’s* approach obscures mvBM’s inaccuracy near the site of the burst, and 3) the accuracy of mvBM depends on the location of the burst in the phylogeny, but Smaers *et al.’s* simulations focus on branches that tend to produce relatively accurate ASRs.

To examine accuracy of ASRs near the site of the burst, we replicated the “burst” simulation conditions used in Smaers *et al.* but varied the location of the burst branch. We generated a pure-birth tree with 100 taxa and allowed each branch to be the “burst branch” for 5 simulations of a continuous trait. The baseline rate of evolution was 0.01, while the burst branch had a rate of 1, replicating the smallest burst size considered in Smaers *et al*. Figure 2 shows the relationship between trait change on the burst branch and error in the ASR at the base of the branch. Reconstructions at the base of the burst branch are strongly biased in the direction of the burst, and the degree of bias depends on the size of the burst. In these simulations, the median distance from the largest to the smallest simulated value is only 0.92, so the errors shown in Figure 2 (ranging from −0.49 to 0.57) are quite large. Because Smaers *et al.’s* accuracy measures are aggregated across the entire tree, their results do not reveal this extreme bias near the site of the burst.

**Figure 2:**
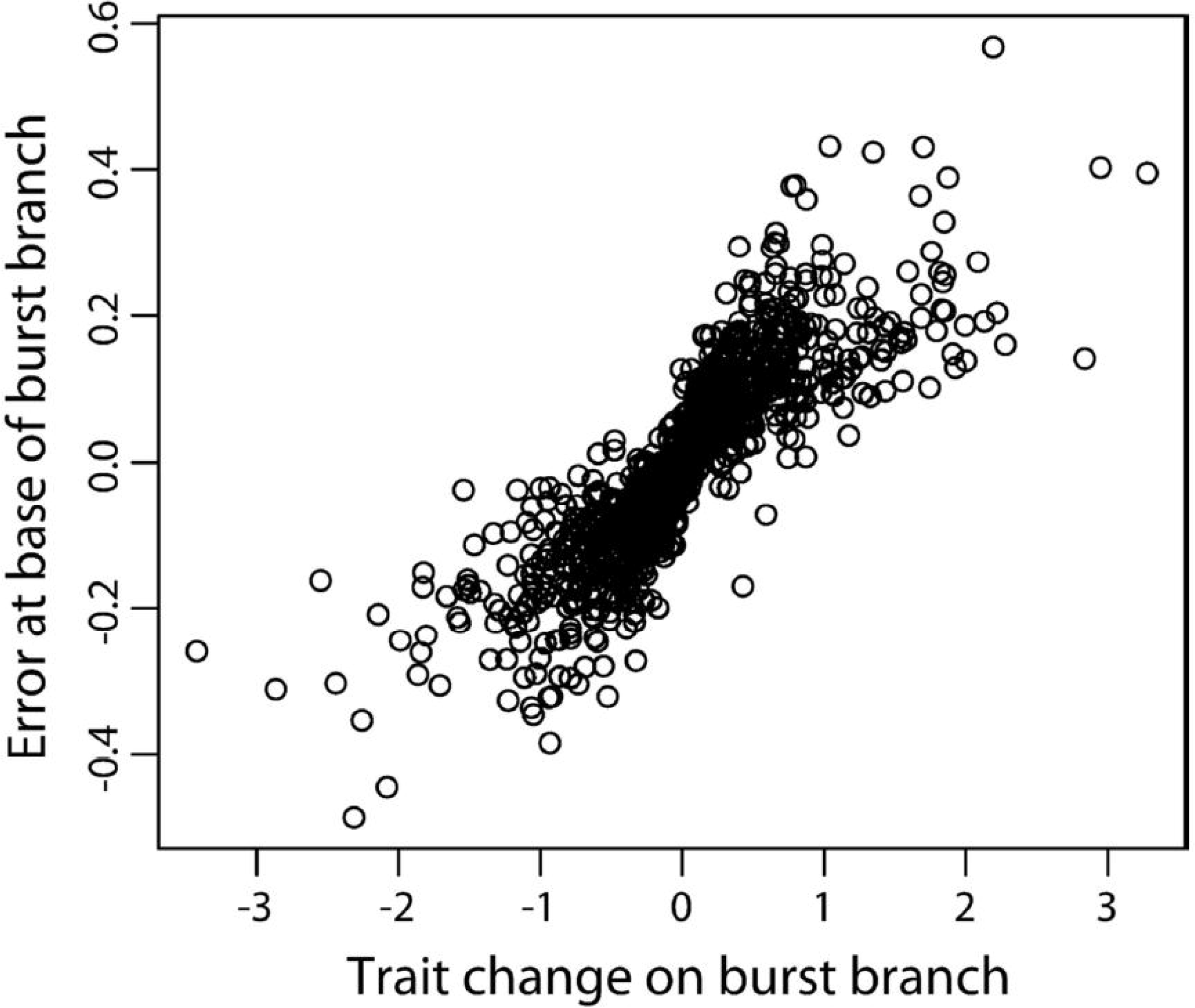
Relationship between error in the estimated ancestral state at the base of the burst branch, and the direction and magnitude of trait change on the burst branch.

Next, we demonstrate that the *R*^2^ of estimated versus true ancestral states is much lower when the number of taxa in the tree decreases. We simulated a pure-birth phylogeny with 30 tips and 5800 continuous traits, allowing each branch to be the burst branch for 100 simulations. We regressed estimated versus true ancestral states for each simulation, as in Smaers *et al*. Figure 3 shows that with the smaller tree, *R*^2^ values are lower and more variable than the results presented by Smaers *et al.* Rather than the *R*^2^ consistently being >0.95, it ranges from nearly 0 to 1, and the median is 0.81 (Figure 3). Smaers *et al.* report higher *R*^2^ values because including more taxa dilutes the effect of the evolutionary burst, resulting in a stronger correlation between estimated and true ancestral states. Thus, the extreme inaccuracy of mvBM ASRs near the site of the burst was obscured by Smaers *et al.’s* use of aggregated accuracy measures across a very large phylogeny relative to the number of bursts.

**Figure 3:**
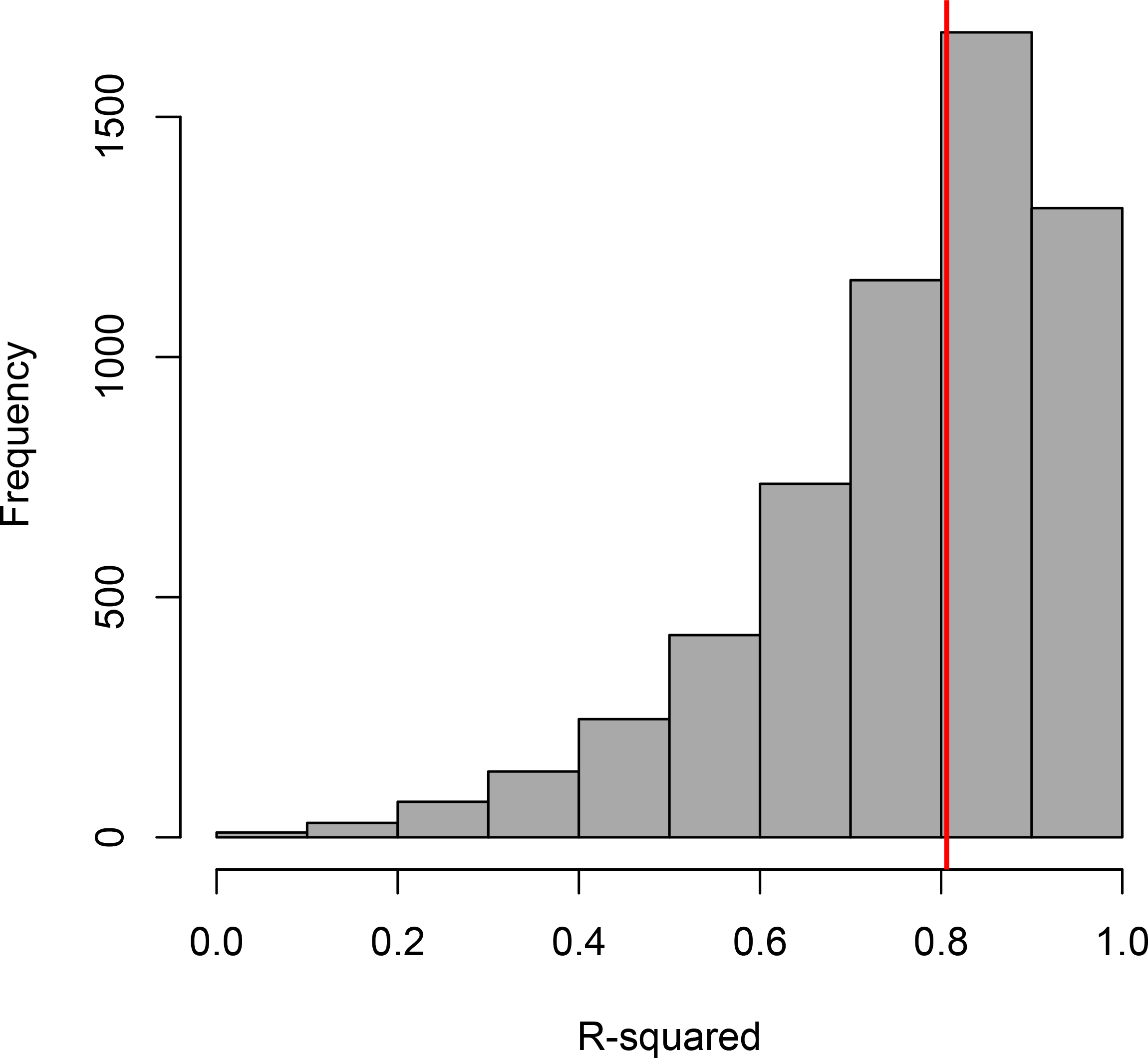
Distribution of *R*^2^ values of estimated versus true ancestral states, given a 30-taxa tree and 5800 simulations in which each of the 58 branches are allowed to be the burst branch for 100 Brownian motion simulations of a continuous trait. The baseline variance of the Brownian motion process is 0.01, while the burst branch has a variance of 1. The red line indicates the median *R^2^*.

Additionally, Figure 4 depicts the median *R*^2^ according to which branch was the burst branch, and illustrates that bursts near the tips yield lower median *R*^2^ values relative to bursts deeper in the tree. This is unsurprising since a burst of evolution will have the strongest impact on nodes ancestral to the burst, such that when the burst occurs near the tips, more ancestral nodes will be affected than when the burst occurs near the root. Because Smaers *et al.* only simulate bursts relatively deep in the tree (their Figure 2), their simulation conditions are favorable toward mvBM.

**Figure 4:**
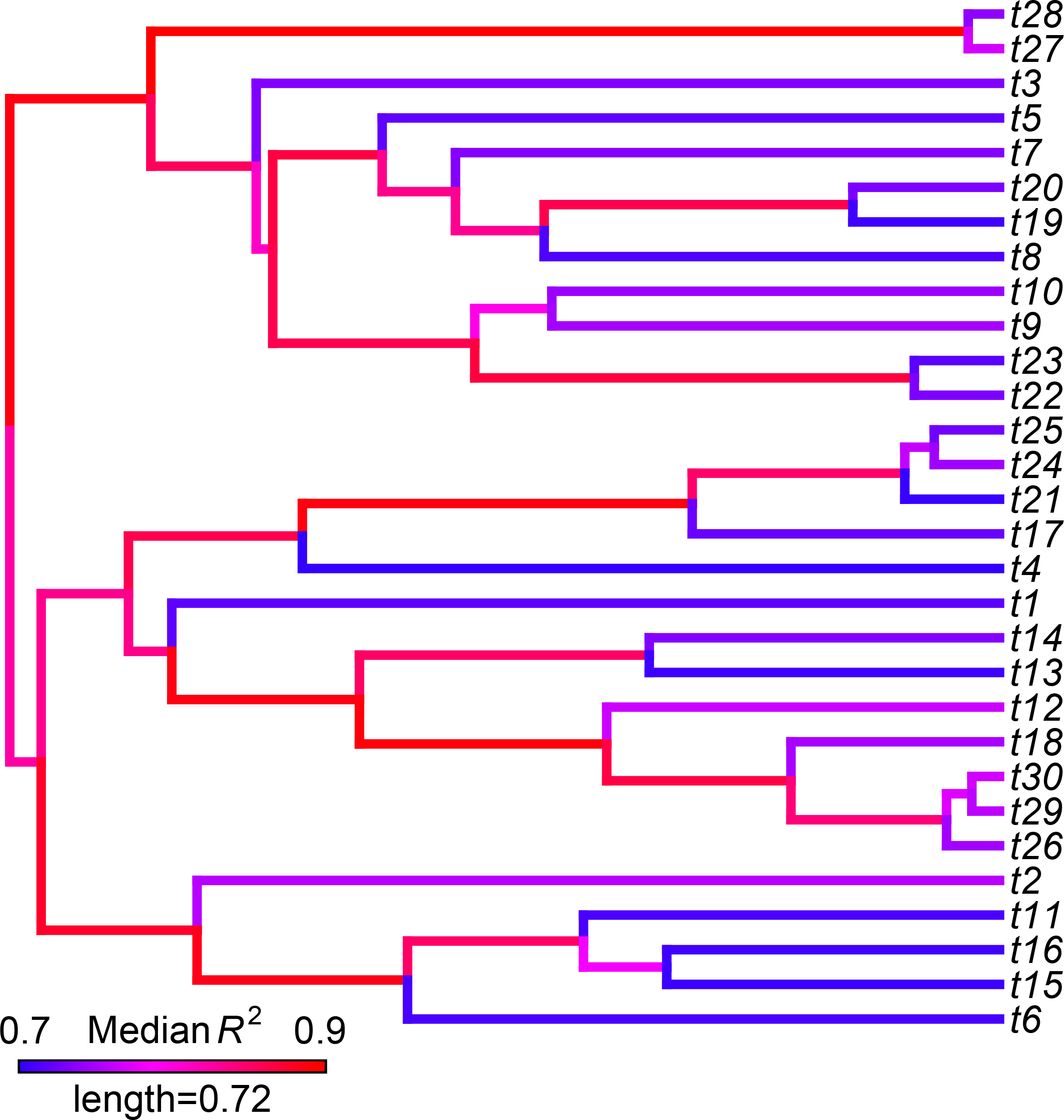
Median *R*^2^ values of estimated versus true ancestral states according to which branch of the phylogenetic tree was the “burst branch”. Branches near the tips have lower *R*^2^ values than branches near the root.

Supplementary Material S3 shows that the five-burst scenario also produces substantially lower *R*^2^ values given a smaller tree and random burst locations.

## Conclusions

Like the IE method (Smaers & Vinicius, 2009), mvBM has substantial theoretical and statistical problems. In contrast to the claims of Smaers *et al.,* mvBM does not include a statistical model of evolution and is not truly a “multiple-rate BM model”. Our primary concern is that the core of mvBM is a circular procedure that overfits phylogenetic branch lengths to data.

We have also demonstrated specific failings in Smaers *et al.*’s simulation study. First, rather than using simulations to test the absolute accuracy of mvBM ASRs, Smaers *et al.* evaluated mvBM relative to BM (a poorly suited method given the simulation conditions) and reported aggregated measures of accuracy rather than accuracy at specific nodes. By reporting results for specific nodes, we demonstrated that mvBM produces extremely biased ASRs near the site of a burst under the exact simulation conditions used by Smaers *et al.* Second, outside of the narrow simulation conditions in Smaers *et al.* (i.e., a large number of taxa relative to bursts, a burst branch deep in the tree), mvBM’s accuracy declines substantially even according to the aggregated measures of accuracy used by Smaers *et al.* We conclude that mvBM is not a viable PCM.

## Acknowledgements

We are indebted to Chris Venditti and Charles Nunn for discussions and thoughtful feedback that improved our manuscript. We also thank. J.D. Pampush for helpful feedback on an early version of the manuscript. Finally, we are grateful to two anonymous reviewers who pushed us to tighten our arguments and address the most fundamental issues with the method in question.

